# A novel deep-sea bacterial threonine dehydratase drives cysteine desulfuration and hydrogen sulfide production

**DOI:** 10.1101/2020.12.23.424250

**Authors:** Ning Ma, Yufan Sun, Wen Zhang, Chaomin Sun

**Affiliations:** CAS Key Laboratory of Experimental Marine Biology, Institute of Oceanology, Chinese Academy of Sciences, Qingdao 266071, China; Laboratory for Marine Biology and Biotechnology, Qingdao National Laboratory for Marine Science and Technology, Qingdao 266071, China; College of Earth Science, University of Chinese Academy of Sciences, Beijing 100049, China; Center for Ocean Mega-Science, Chinese Academy of Sciences, Qingdao, 266071, China; Department of Medical Microbiology, Key Laboratory of Medical Molecular Virology of Ministries of Education and Health, School of Basic Medical Sciences, Fudan University, Shanghai 200032, China; Fudan University Pudong Medical Center, Shanghai Key Laboratory of Medical Epigenetics, Institutes of Biomedical Sciences, Fudan University, Shanghai 200032, China; The Department of Systems Biology for Medicine, School of Basic Medical Sciences, Fudan University, Shanghai 200032, China

## Abstract

Cysteine desulfuration is one of the main ways for hydrogen sulfide (H_2_S) generation in cells and is usually conducted by cystathionine γ-lyase. Herein, we describe a newly discovered deep-sea bacterial threonine dehydratase (psTD), which is surprisingly discovered to drive L-cysteine desulfuration. The mechanisms of psTD catalyzing cysteine desulfuration towards H_2_S production are first clarified *in vitro* and *in vivo* through a combination of genetic and biochemical methods. Furthermore, based on the solved structures of psTD and its various mutants, two or three pockets are found in the active site of psTD, and switch states between inward and outward orientation of a key amino acid R77 determine the open or close status of Pocket III for small molecule exchanges, which further facilitates cysteine desulfuration. Our results reveal the functional diversity and structural specificity of psTD towards L-cysteine desulfuration and H_2_S formation. Given the broad distribution of psTD homologs in different bacteria, we speculate that some threonine dehydratases have evolved a novel function towards cysteine desulfuration, which benefits the producer to utilize cysteine as a sulfur source for better adapting external environments.

## INTRODUCTION

The important life-supporting role of hydrogen sulfide (H_2_S) has been found from bacteria to plants, and finally to mammals (Wang, 2012). In bacteria, endogenous H_2_S is involved in stress responses, such as oxidative stress and antibiotics, and the assembly of intracellular [Fe-S] clusters which are ubiquitous and evolutionary ancient prosthetic groups required to sustain fundamental life processes (Johnson et al, 2005; Mihara & Esaki, 2002; Mironov et al, 2017; Shatalin et al, 2011). L-cysteine is a common substrate in many bacterial species for H_2_S generation, which is catalyzed by cysteine desulfurases using pyridoxal 5’-phosphate (PLP)-based chemistry (Chiku et al, 2009; Mihara & Esaki, 2002; Szabo, 2018; Wendisch, 2007). Bacterial cysteine desulfurases include cystathionine β-synthase (CBS), cystathionine γ-lyases (CSE), *O*-acetylserine sulfhydrylase (OASS), et al (Awano et al, 2005; Devi et al, 2017; Dunleavy et al, 2016). In *Escherichia coli*, five different enzymes have been identified to possess cysteine desulfurase activity, which including cystathionine β-lyase (MetC), cysteine synthase A/*O*-acetylserine sulfhydrylase A (CysK), cysteine synthase B/*O*-acetylserine sulfhydrylase B (CysM), β-cystathionase (MalY), and tryptophanase (TNaA) (Awano et al, 2005).

PLP-dependent enzymes catalyze manifold reactions of amino acid metabolism (Alexander et al, 1994). Threonine dehydratase (TD), also named threonine deaminase, belongs to the β-family of PLP-dependent enzyme catalyzing the formation of α-ketobutyrate and NH_3_ from L-threonine (Schomburg & Salzmann, 1990; Simanshu et al, 2006). It also catalyzes the deamination of L-serine, L-homoserine, β-chlora-L-alanine and L-allothreonine (Schomburg & Salzmann, 1990). Two types of TD have been found in bacteria: the biosynthetic threonine dehydratase (BTD) and the catabolic threonine dehydratase (CTD) (Yu et al, 2013). BTD, encoded by the gene *ilvA*, is expressed under aerobic conditions and catalyzes the first reaction in the isoleucine biosynthesis pathway (Gallagher et al, 1998). L-isoleucine and L-valine act as an allosteric inhibitor and an activator, respectively (Gallagher et al, 1998). IlvA protein provided one of the earliest examples of feedback inhibition (Umbarger, 1956). CTD, encoded by the gene *tdcB*, is induced anaerobically and catalyzes the first reaction in the degradation of L-threonine to propionate (Simanshu et al, 2007). Unlike IlvA, TdcB protein is insensitive to L-isoleucine and L-valine and is activated by AMP (Simanshu et al, 2006). In *E. coli*, cysteine has been shown to inhibit TD activity, resulting in transient amino acid starvation, which can be reversed by threonine (Harris, 1981). However, TD has not previously been shown to have a cysteine desulfurase activity.

In the current work, a deep-sea bacterium *Pseudomonas stutzeri* 273 was found to produce substantial amounts of H_2_S in the presence of cysteine, and TD of this bacterium (psTD) was demonstrated to drive L-cysteine desulfuration. A combination of the proteomic method together with gene knockout approaches revealed that psTD drove L-cysteine desulfuration and thereby the generation of H_2_S in *P. stutzeri* 273. Furthermore, structural insights of psTD mediating L-cysteine desulfuration were detailedly disclosed. Overall, this work reports a previously undocumented cysteine desulfurase activity for bacterial TD both *in vivo* and *in vitro*.

## RESULTS

### psTD drives L-cysteine desulfuration *in vivo*

Originally, we observed that addition of L-cysteine promoted *P. stutzeri* 273 to generate substantial amounts of H_2_S (Fig. 1A). To investigate the response of *P. stutzeri* 273 when exposed to L-cysteine, we performed proteomic analyses of *P. stutzeri* 273 incubated in LB and LB supplemented with 8 mmol/L L-cysteine. Among the significantly up-regulated proteins, four were associated with sulfur metabolism (Fig. 1B). Meanwhile, the expression of pyridoxal kinase for pyridoxal 5’-phosphate (PLP) synthesis was significantly increased (Fig. 1B). Of the four proteins, psTD is one of the proteins containing a PLP binding domain (Fig. S1), which is an obligate cofactor required for cysteine desulfuration (Majtan et al, 2018; Mihara & Esaki, 2002; Sun et al, 2009). Based on the proteomic results, we propose that psTD might involve in the cysteine metabolism of *P. stutzeri* 273.

**Fig. 1.**
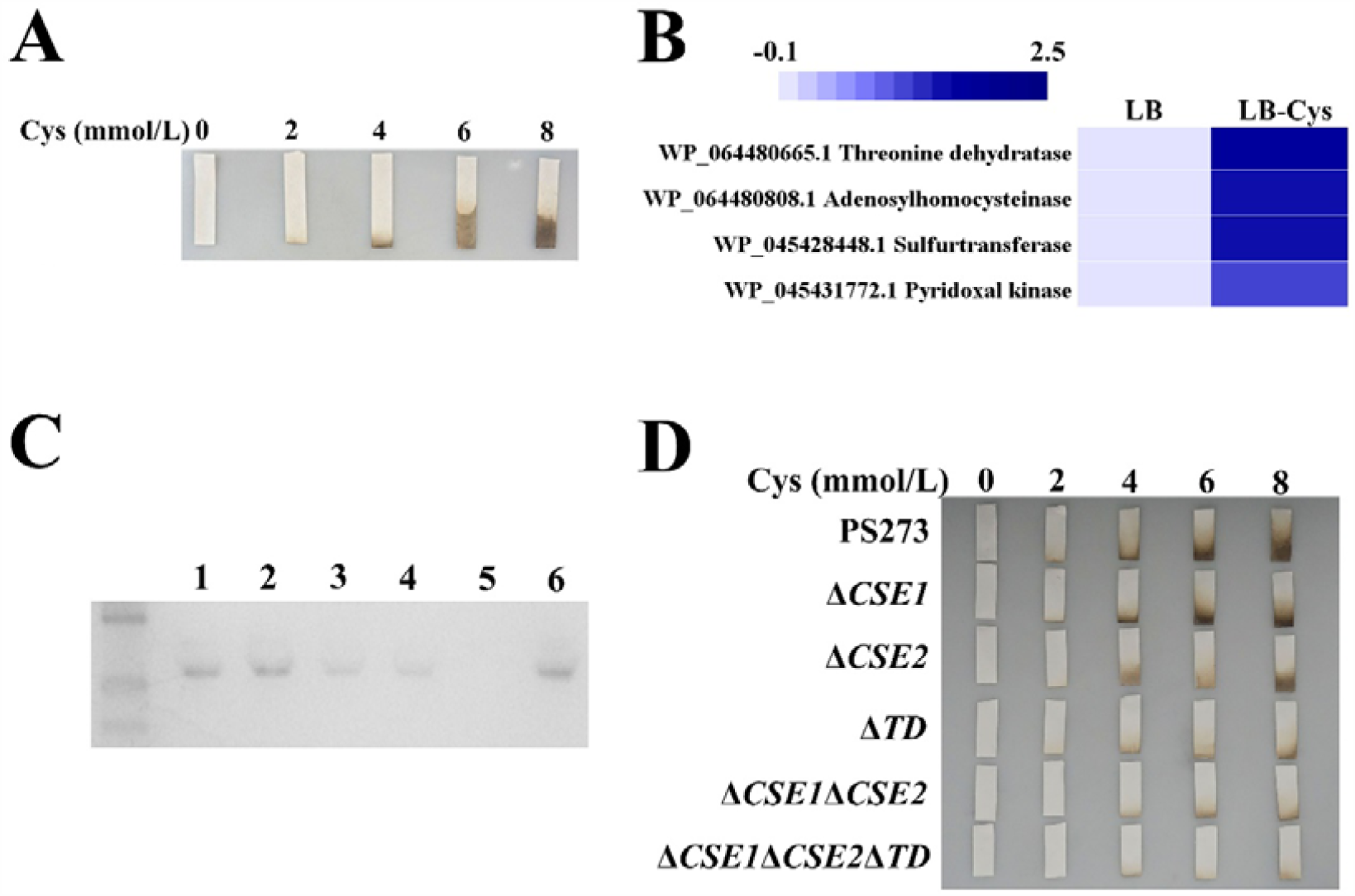
Key roles of psTD in driving intracellular H_2_S generation through degrading L-cysteine in *P. stutzeri* 273. (**A**) H_2_S production of *P. stutzeri* 273 challenged with different concentrations of Cys. (**B**) Proteomic assay of the relative level of proteins associated with cysteine metabolism in *P. stutzeri* 273 under LB alone and LB with 8 mmol/L L-cysteine. (**C**) The in-gel activity assay with bismuth staining applied for detecting the proteins that produced H_2_S from L-cysteine in *P. stutzeri* 273 wild type (PS273), mutant strains (Δ*CSE1*, Δ*CSE2*, Δ*CSE1*Δ*CSE2*, Δ*TD*), and the psTD encoding gene complementary strain Δ*TD*/c*TD*. Lane 1: PS273, Lane 2: Δ*CSE1*, Lane 3: Δ*CSE2*, Lane 4: Δ*CSE1*Δ*CSE2*, Lane 5: Δ*TD*, Lane 6: Δ*TD*/c*TD*. (**D**) H_2_S production of PS273 and its mutant strains incubated in LB supplemented with different concentrations of Cys for 12 h.

To evaluate corresponding roles of psTD and other two cystathionine γ-lyases determining the production of H_2_S in *P. stutzeri* 273, we constructed a series of deletion mutants targeting the genes encoding psTD (Δ*TD*), cystathionine γ-lyase 1 (Δ*CSE1*), cystathionine γ-lyase 2 (Δ*CSE2*), or different combinations. Thereafter, cell extracts from wild-type *P. stutzeri* 273 and the corresponding mutants were analyzed using in-gel activity assays for detecting the potential H_2_S-producing enzymes. Clearly, wild-type *P. stutzeri* 273 could degrade L-cysteine to produce H_2_S, as could the mutants Δ*CSE1*, Δ*CSE2*, and Δ*CSE1*Δ*CSE2* (Fig. 1C). However, the ability to generate H_2_S was compromised in Δ*TD* (Fig. 1C), which showed lower levels of H_2_S production than wild-type *P. stutzeri* 273 (PS273), Δ*CSE1*, Δ*CSE2*, and Δ*CSE1*Δ*CSE2* (Fig. 1D). Complementing Δ*TD* with the wild-type psTD gene (Δ*TD*/c*TD*) restored H_2_S production (Fig. 1C). Thus, we conclude psTD plays a key role in catalyzing L-cysteine desulfuration and H_2_S formation in *P. stutzeri* 273 *in vivo*.

### psTD drives L-cysteine desulfuration *in vitro*

To determine whether psTD was capable of catalyzing H_2_S generation from L-cysteine *in vitro*, it was over-expressed and purified from the *E. coli* BL21 cell line. Recombinant psTD showed a strong absorbance at 412 nm, a typical symbol of covalently bound PLP (Fig. 2A), and it is consistent with the presence of a PLP binding domain in psTD (Fig. S1). psTD generated H_2_S from L-cysteine in the in-gel activity assay (Fig. 2A), consistent with the *in vivo* assay (Fig. 1C). Moreover, we identified L-serine as one of the downstream products (Fig. 2B) based on a previous report (Chiku et al, 2009). Pyruvate, a downstream product of L-serine, was also detected by spectrophotometry (Fig. 2C). In addition, psTD had a more substantial catalytic velocity when compared to other CSEs according to the intrinsic kinetics of the reaction (Table S2). Together, we propose that psTD catalyzes the following reaction: L-cysteine + H_2_O → L-serine + H_2_S → pyruvate + NH_3_ + H_2_S (Fig. 2D).

**Fig. 2.**
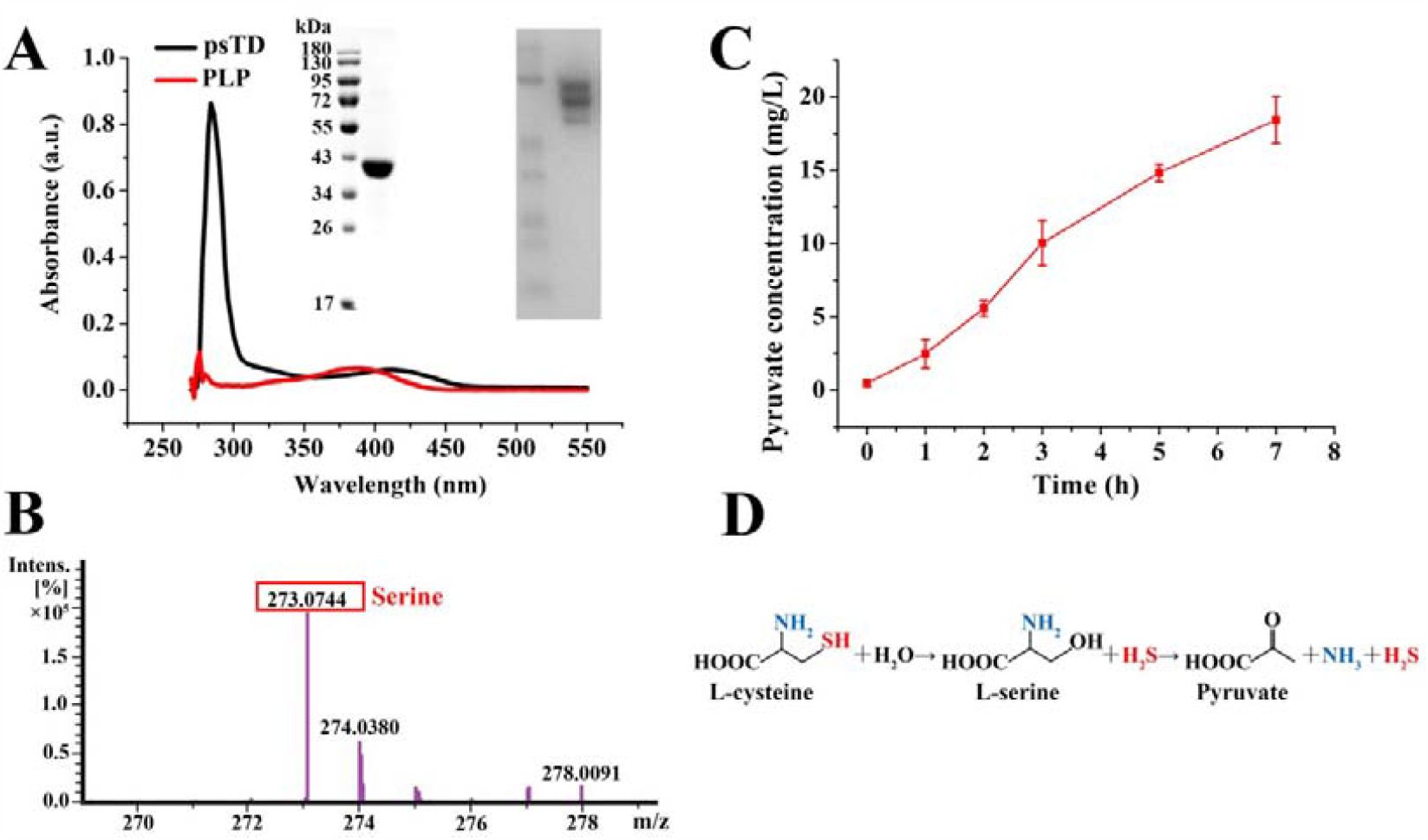
Products determination of L-cysteine catalyzed by psTD. (**A**) Absorbance spectrum of purified psTD in Tris buffer (50 mmol/L Tris, 150 mmol/L NaCl, 250 mmol/L imidazole, pH 8.0) (black line) and unbound pyridoxal 5’-phosphate (PLP) (0.02 mmol/L in Tris buffer, red line). Insert in panel A: SDS-PAGE gel (left) of purified psTD (20 μg) showing a size around 40 kDa and Native-PAGE gel (right) with bismuth staining for 30 min showing psTD (0.4 μg) with the ability of catalyzing H_2_S generation from L-cysteine. (**B**) LC-MS analysis of the reaction products catalyzed by psTD. (**C**) Determination of pyruvate concentration in the enzymatic mixture catalyzed by psTD. (**D**) The proposed reaction of L-cysteine catalyzed by psTD.

### Structural basis of psTD catalyzing L-cysteine desulfuration

To better understand the catalytic mechanism of psTD-driven L-cysteine desulfuration, the crystal structures of psTD wild type and corresponding mutant, and complex bound with PLP were determined (Table 1). The crystal structures of psTD were solved around 1.5∼2Å in two conditions, C6 (PDB 7DAP) and E11 (with high concentration NH_4_^+^, PDB 7DAQ). The structure of psTD is an asymmetric homodimer possessing two similar subunits (superposition with RMSD=0.20Å) (Fig. S2A). Each subunit is highly conserved to the structure of 1TDJ (TD from *E coli*, superposition with RMSD=1.67Å), although the identity is only 31% between two sequences (Fig. S2B).

**Table 1.**
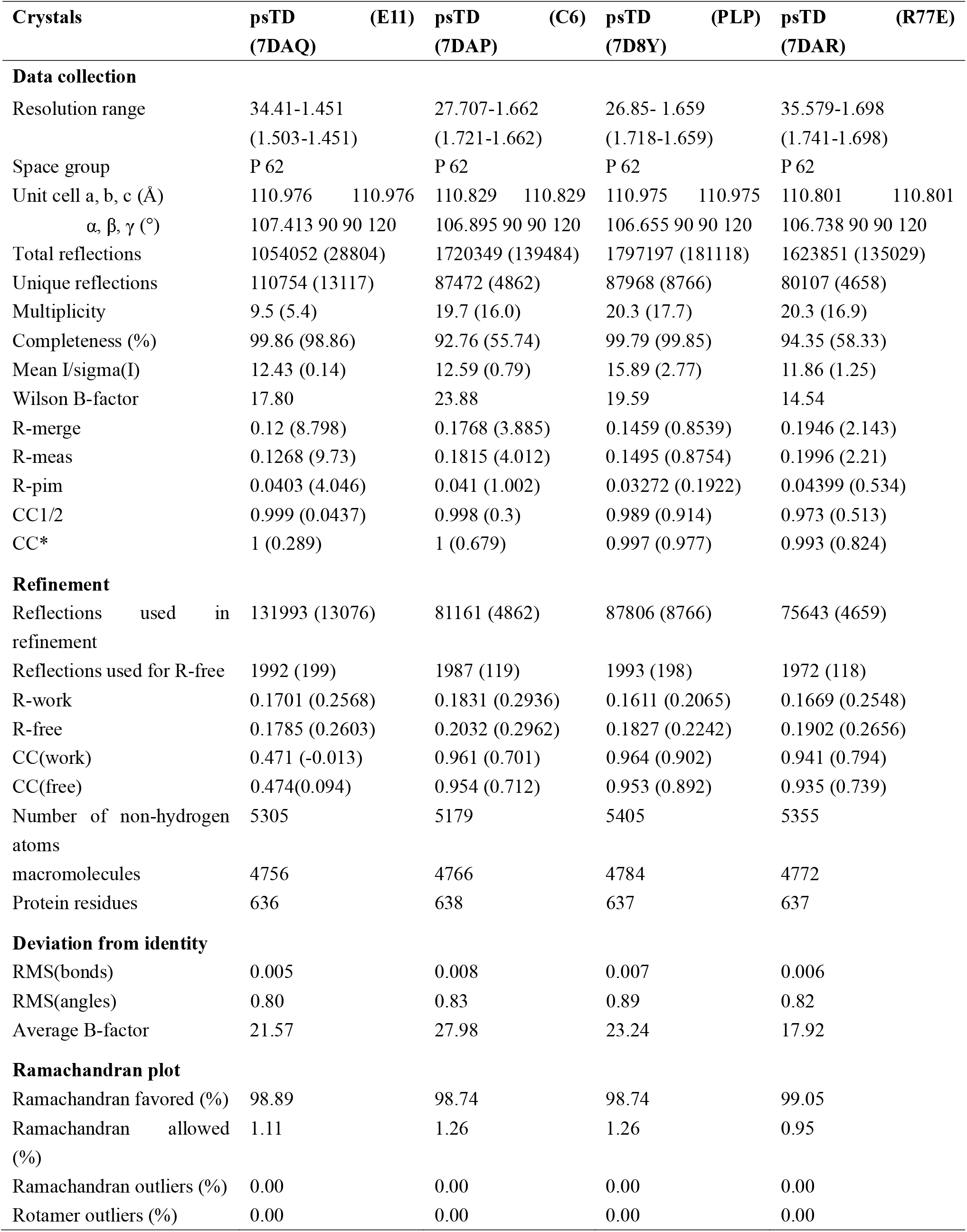
Diffraction data and refinement statistics of psTD, psTD/PLP and psTD (R77E).

As PLP is a co-enzyme of TD and is essential for TD to perform catalytic activity, thus co-crystal structure of complex containing psTD and PLP was solved and analyzed (PDB 7D8Y). The structural results show that PLP binds to the amino acid K51 with both covalent and non-covalent bonds and occupies one pocket of active site from several resolved crystals (Fig. 3A). Consistently, the sequence alignment of TD protein families indicates that the key residue of active site is K51 (Fig. S3A). It is noting that PLP may be missing in the site when high concentration NH_4_^+^ is present in the crystallization buffer since NH_4_^+^ may occupy one or two pockets around K51 in the active site (Fig. 3B). Moreover, the sidechain of the amino acid R77 interacts with PLP in C6 crystal structure while it is orientating to protein surface in high NH_4_^+^ crystal structure since the active site is occupied by NH_4_^+^ (Fig. 3C). Therefore, NH_4_^+^ may prevent the sidechain of R77 from orientating inside of protein as well as binding with PLP, which is consistent with the proposal that NH_4_^+^ (as a catalytic product) may reduce substrate concentration and slow down the reaction as negative feedback (Figs. 2D and 3B). The reduced activity of psTD caused by NH_4_^+^ further confirms this view (Fig. 3D). Interestingly, the conformation change and alternative conformation of R77 sidechain observed in chain A and chain B imply the dynamics and specific function of R77 (Fig. 3C and Fig. S4A). To explore the potential function of R77, the structure of psTD with mutation of R77E (R is mutated to E; PDB 7DAR) was also solved for comparison. The results show that there are alternative sidechains for E77 in chain A and alternative mainchain for amino acid G78 in chain B of 7DAR, where the sidechain of amino acid E77 is extending to protein surface (Fig. 3E).

**Fig. 3.**
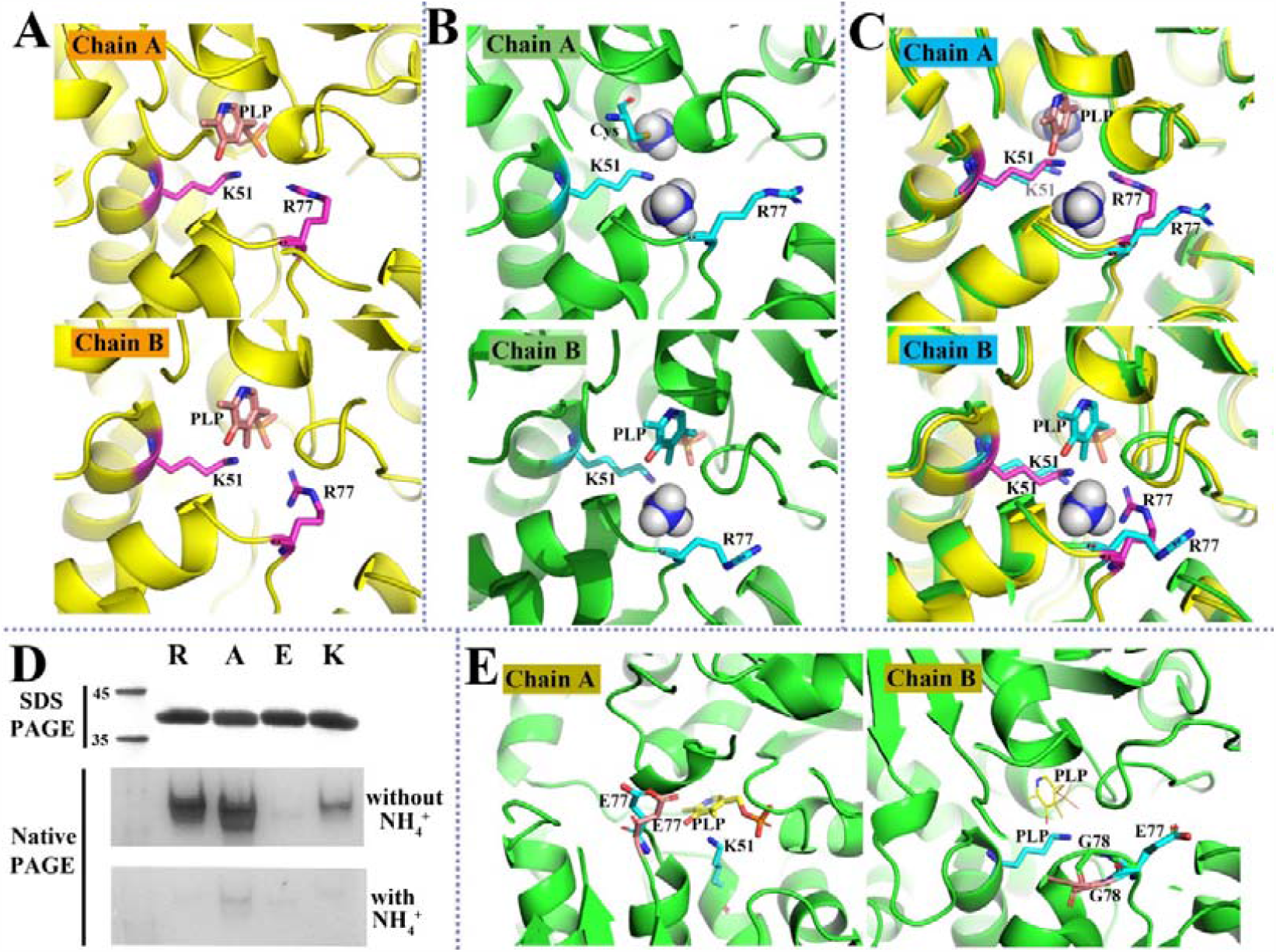
Structure comparison of different psTD crystal structures. (**A**) PLP bound to K51 with possible covalent bond and covalent bond in chain A and chain B of structure 7DAP (carbon in purple for key residues), respectively. (**B**) An amino acid (e.g. cysteine) bound to K51 in chain A of structure 7DAQ (two NH_4_^+^, carbon in cyan for key residues); PLP bound to K51 with non-covalent bond in chain B of 7DAQ (one NH_4_^+^, R77 extending orientating outside for both chain A and B). (**C**) Structure comparison of 7DAQ and 7DAP for chain A (one NH_4_^+^ overlapping with PO_4_^-^ of the virtually corresponding PLP) and for chain B. (**D**) SDS-PAGE gel and Native-PAGE gel of purified psTD wild type (R) and variants R77A (A), R77E (E), and R77K (K) with bismuth staining (without or with 2 mol/L NH_4_^+^) for 5 min. The loading amount of each lane is 9 μg. (**E**) Dual conformation of E77 in chain A (7DAR, carbon in cyan for key residues, carbon in orange for alternative E77); Dual conformation G78 in chain B (7DAR, with the virtually corresponding PLP, carbon in orange for alternative G78).

The inside active sites in different psTD structures were further compared by the map of electrostatic potential (MEP). PLP binds deep inside the psTD structure with positive charged pockets around catalytic key residue K51 (Fig. 4A). Besides, the active site of psTD is unique in sharp and could be switchable at the gate position near R77 in some structure states (Fig. 4B). In the structure 7DAQ, there are an amino acid residue bound by K51 and two NH_4_^+^ ions instead of PLP in chain A, while PLP with weak density is present in chain B with non-covalent binding to K51, together with one NH_4_^+^ ion in another pocket (Figs. 3B and 4B). In the structure 7DAR, the E77 with negative charged sidechain may block the gate of the active site by negative potential gap at the psTD surface (Fig. 4C). Consistently, psTD with mutation R77E significantly reduces the catalytic activity to form H_2_S (Fig. 3D).

**Fig. 4.**
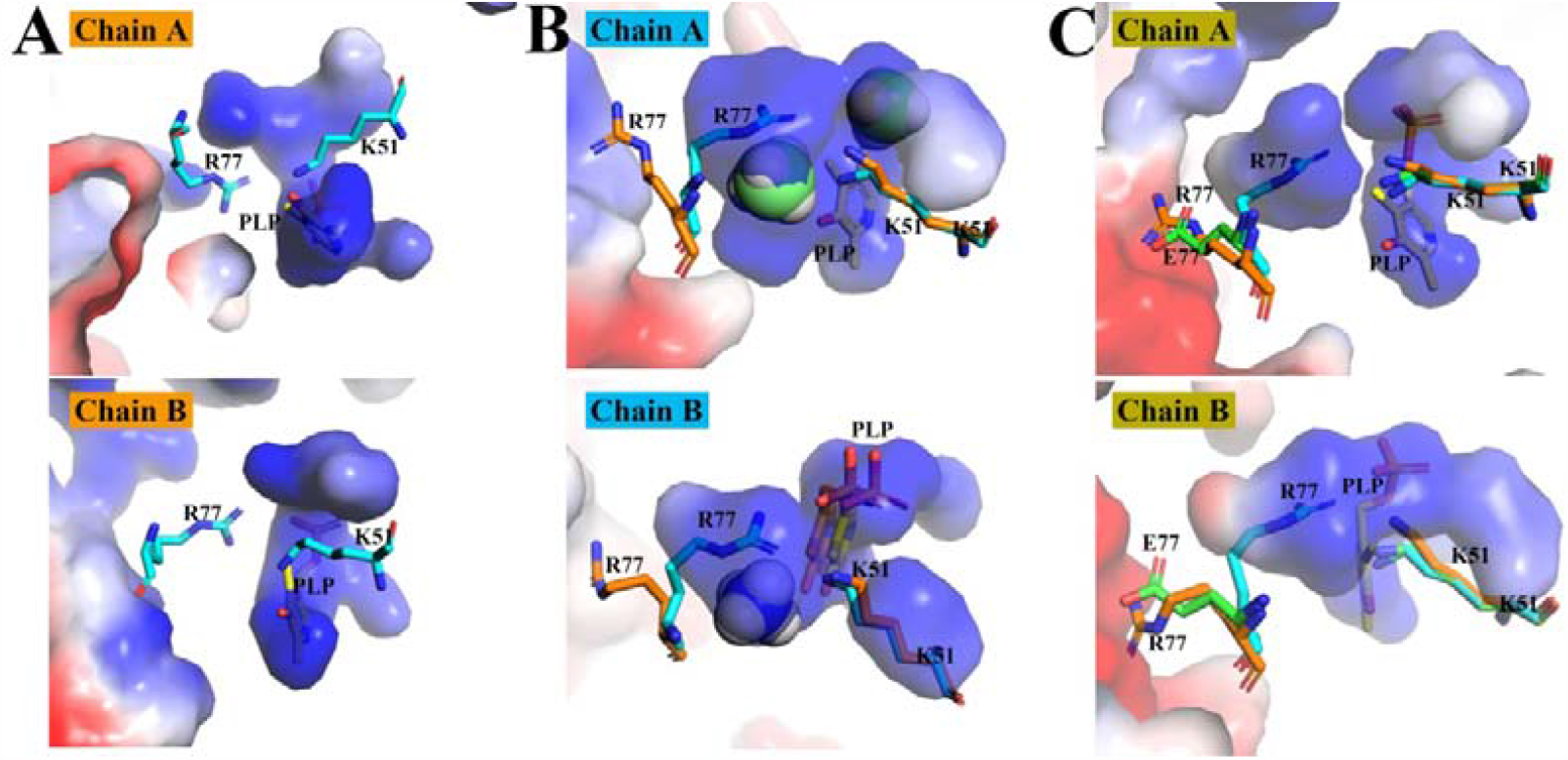
The map of electrostatic potential (MEP) comparison of the inside active sites from different psTD structures. (**A**) MEP inside of structure 7DAP (chain A and B, carbon in cyan). (**B**) MEP of 7DAQ and key residues comparison for 7DAQ (carbon in orange) and 7DAP (chain A and B). (**C**) MEP of 7DAR and key residues comparison for 7DAR (carbon in green), 7DAP and 7DAQ (chain A and B).

Furthermore, there are intensive hydrogen-bond interactions between phosphate group of PLP and amino acids G177-G181 (GLGSG), A277, S302 as well as the salt bridge interaction between phosphate group of PLP and the specific residue R77, therefore these residues may stabilize the PLP binding (Fig. 5). Notably, the sequence motifs around amino acids F50-K51, A75-H80 and G177-G181 are conserved through whole TD family (Fig. 5E), indicating the threonine dehydratase function of psTD is conserved and ubiquitously exists in different microorganisms. While the unique cysteine desulfuration may result from specific structure and motifs associated with the diversity positions, such as R77 and the relevant residues near active site.

**Fig. 5.**
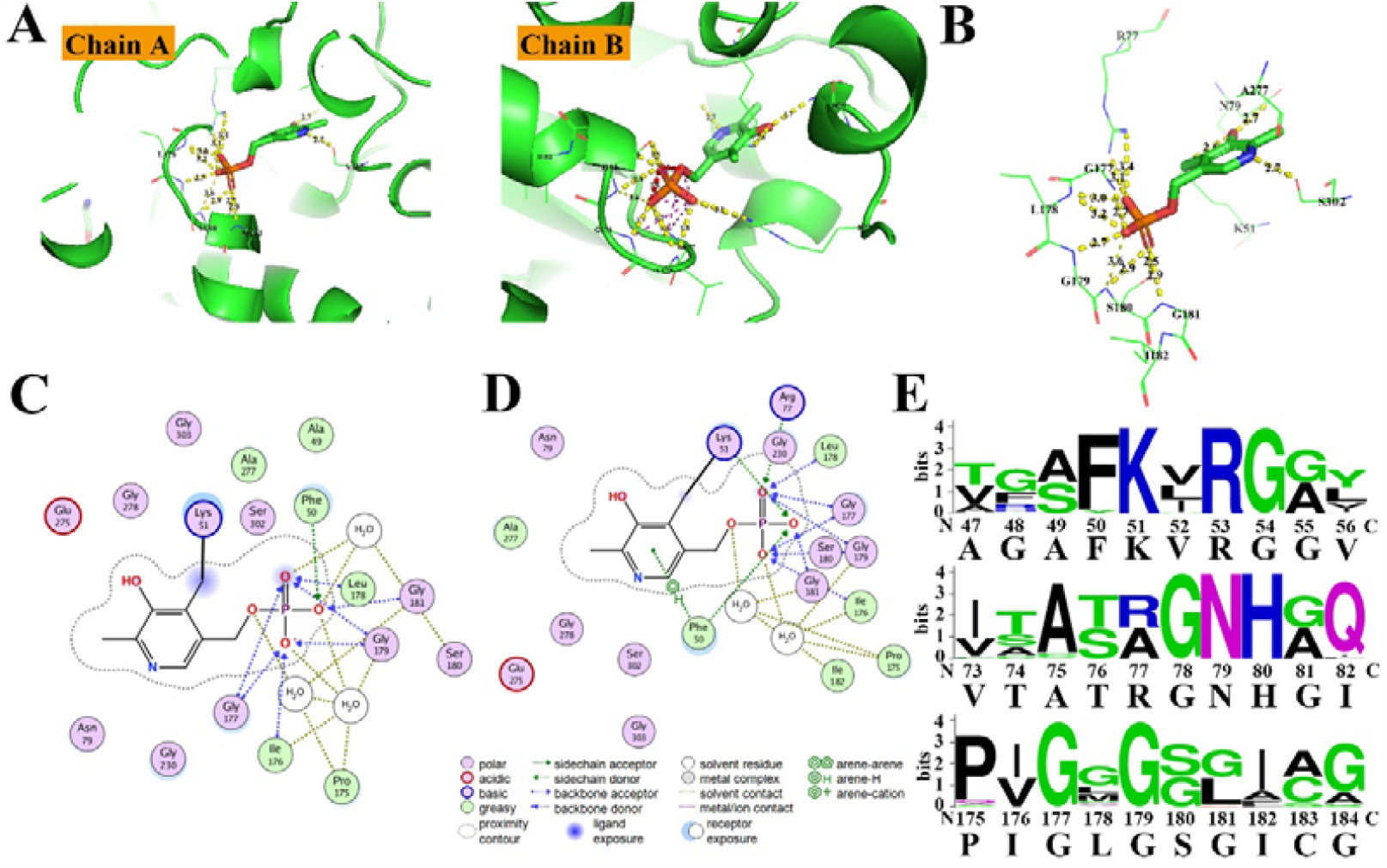
The molecular interactions between PLP and psTD near the active site. (**A**) The polar interactions between PLP and psTD structure (chain A and B of 7D8Y, the same as below, dashing line indicates the possible hydrogen bonds). (**B**) The PLP contacting residues of psTD (chain A). (**C, D**) 2D plot of the interactions between PLP and psTD (chain A and B). (**E**) Weblogo for the amino acids around K51, R77 and G177-G181. The size of a single letter amino acid code in the sequence logo represents the occurrence of a particular amino acid at a particular position. The numbers are the locations of amino acids in psTD. psTD sequence is underneath of the horizontal axis.

### Two or three pockets in the active site determine the catalytic function of psTD

Notably, MEP comparison of active sites between structures 7D8Y (psTD-PLP complex) and 1TDJ (TD homolog from *E. coli*) shows that the active site for 7D8Y is close to U shaped channel which could be divided into two or three pockets (Fig. 6 and Fig. 7). Pocket I (PI) is occupied by PLP and Pocket II (PII) is on another side of K51 (Figs. 7A and 7B). Pocket III (PIII), the access channel, opens at the surface of 7D8Y around R77 and only presents with R77 outward orientation (Figs. 6, 7C, and 7D). The residues interacting with PLP (as PI of active site) are highly conserved at the structural level except flexibility of R77 and G181, although the sequence motifs are relatively similar, together with some diversity positions at T76, R77, L178, S180, G181, A277, S302, et al (Fig. S5B). The structural conservation is consistent with the phylogenetic classification and sequence motifs of TD family (Figs. S2E, S2F, S3B, S7). Interestingly, there is only one pocket to hold PLP at the active sites from 1TDJ, 1P5J, 6VJU (1P5J: Ser dehydratase, 6VJU: Cys synthase), which is directly opened to protein surface (corresponding to PIII) and mostly conserved with PI of 7D8Y at the structural level (Fig. 6). The small molecules of reaction products are favorable to bind in PII, which may result in multiple functions including cysteine desulfuration based on its structure and sequence specificity. Besides the structure difference with pockets, there is also some sequence diversity at L151, C233 for PII and PIII, respectively (Fig. S5).

**Fig. 6.**
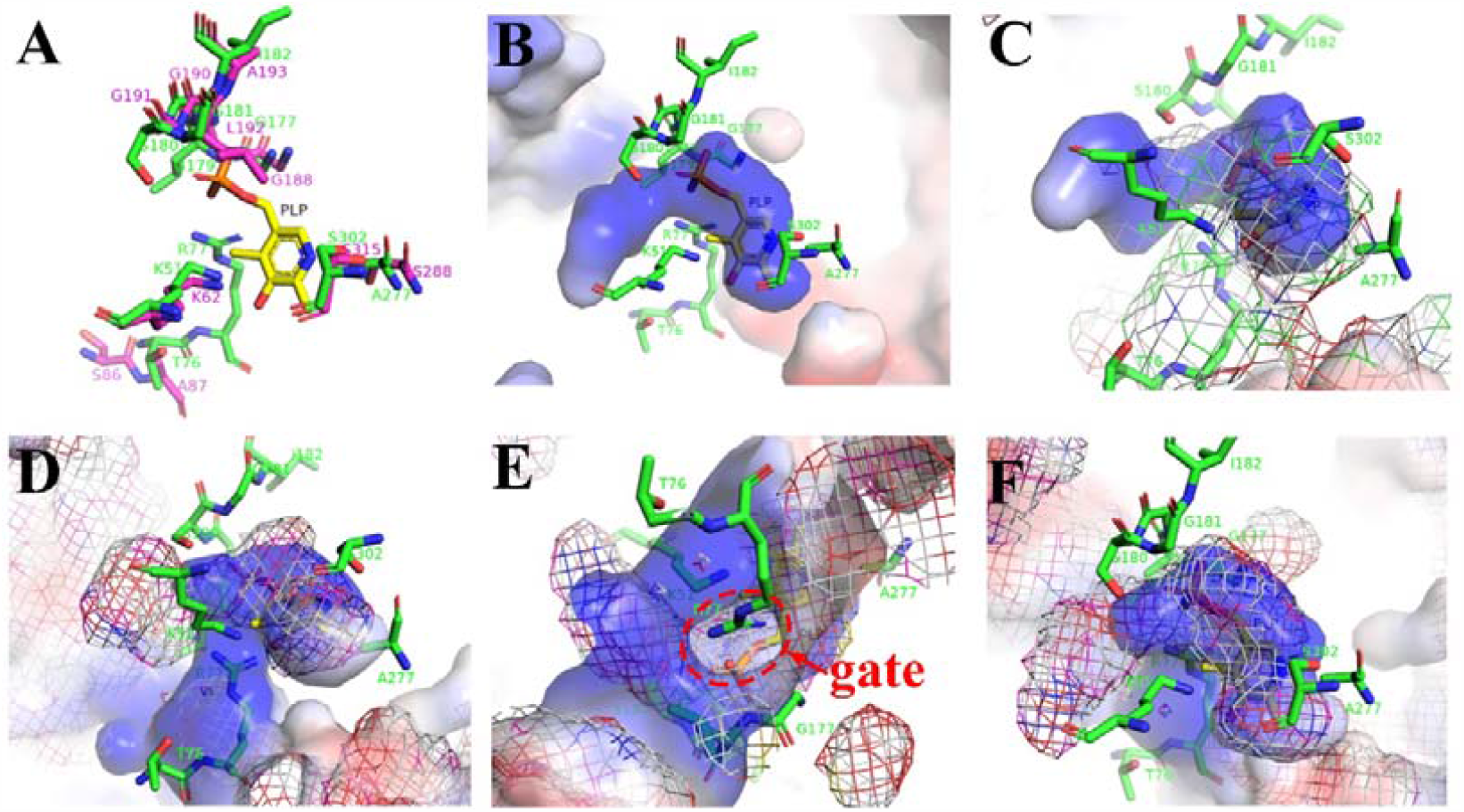
The detailed comparison of active sites from psTD and 1TDJ. (**A**) Key residues comparison around PLP between psTD (chain B of 7D8Y, green carbon) and 1TDJ (purple carbon). (**B**) Pockets of active site from psTD with surrounding residues interacting PLP. (**C**) MEP of psTD (surface) and 1TDJ (mesh). (**D**), (**E**), (**F**) MEP of psTD (mesh) and 1TDJ (surface) from three different views. The access channel (or gate) for active site of 1TDJ is circled with red dashline in (**E**).

**Fig. 7.**
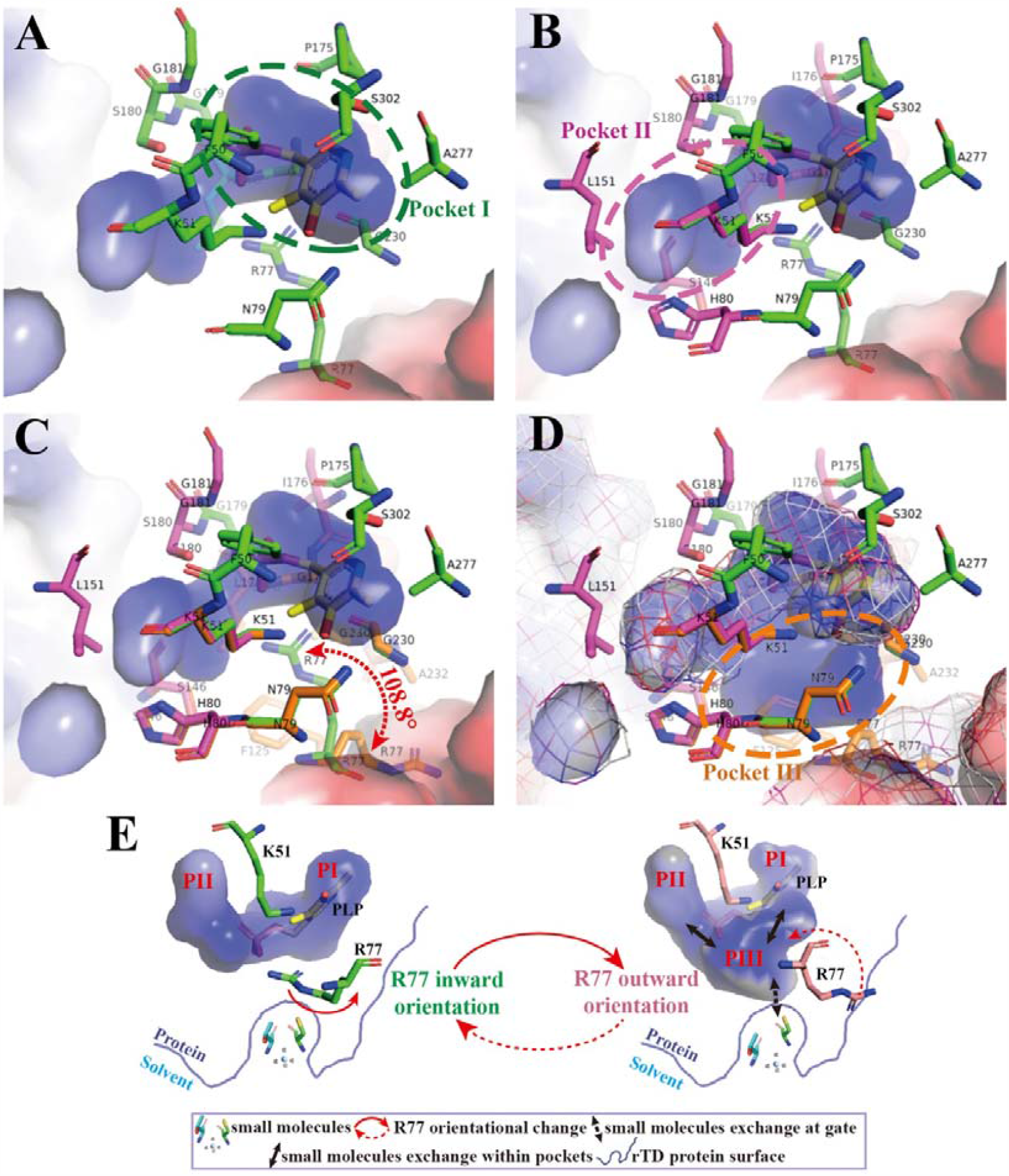
Three pockets of active site of psTD structure. Pocket I (occupied by PLP), Pocket II (for reaction products/small molecules), and Pocket III (access channel) of active site from psTD (chain B of 7D8Y). (**A**) Key residues interacting with PLP at Pocket I (green circle, residues in green carbon). (**B**) Key residues of Pocket I and II (deep purple circle, residues in deep purple carbon). Key residues of Pocket I and II (R77 inward orientation) (**C**), and together with Pocket III (brown circle, residues in brown carbon) (R77 outward orientation), comparing with Pockets I and II MEP in mesh (**D**). Notes: Pocket I and II with R77 inward orientation shown as MEP surface in A, B and C; R77 outward orientation shown in D; both R77 inward and outward conformations compared and shown rotation angle in C. (**E**) Proposed mechanisms of structure dynamics and specificity for psTD enzyme activities: R77 switch states between inward and outward orientation, which results in PIII open or close for small molecule exchange and corresponding reactions.

As mentioned above, the R77 has alternative sidechain conformations in 7D8Y: one is interacting with PLP (inward state with PI and PII), and another is orienting to protein surface (outward state with additional PIII) (Fig. 7E). Active site could expand more space between pockets and protein surface, so that small molecules could access the gate of the active site and reach PLP to trigger the further reactions for R77 outward conformation (Fig. 7). The psTD with mutation of R77A could remain comparable enzyme activity since the gate of active site may keep opened like 1TDJ (with small sidechain of Ala at corresponding position), while psTD with the mutation R77E loses its function since its negative charge may block the small molecules to access the active site (Figs. 3D, 6E, S5A). Therefore, residues near R77 may play a role as gate-keepers controlling the entry to and from the active site, which has shown some extended flexibility of the sidechain of R77 and the mainchain near R77 (Pravda et al, 2014), while the function and mechanism for dynamics of PIII may need further study.

## DISCUSSION

In the present study, detailed biochemical and structural mechanisms of psTD driving L-cysteine desulfuration are first disclosed. The initial clue of psTD mediating L-cysteine desulfuration comes from the proteomic and genetic results (Fig. 1), which propose psTD as a new candidate for metabolizing L-cysteine in the deep-sea bacterium *P. stutzeri* 273 (Fig. 1). TD is principally known for catalyzing L-threonine or L-serine deamination in microorganisms (Ernst & Downs, 2018; Favrot et al, 2018; Lambrecht et al, 2013). In this study, threonine dehydrase was identified as a new member of cysteine desulfurases, and the alternative activities of enzymes play an important role in the diversification of enzymes (O’Brien & Herschlag, 1999). We speculate cysteine desulfuration catalyzed by psTD might contribute to the organic/inorganic sulfur of the deep-sea sediment. Overall, the discovery of cysteine desulfurase activity of psTD introduced a new metabolic pathway for the H_2_S enzymatic production (Wang, 2012).

With biochemical method, the basic process of cysteine desulfuration mediated by psTD is clear (Figs. 2B-2D), however, the deep catalyzing mechanisms are still obscure. With this, we further investigated the underlying mechanisms by structural methods. Based on the structure of psTD, its mutant and complex with PLP, we clarify that the active site with positive charge can be recognized as two or three major pockets for PLP (PI) and substrates (PII) binding or the small molecules exchange (PIII), which are essential in cysteine desulfuration mediated by psTD. PII and PIII may play a key role in helping determine enzyme specific substrates (cysteine and serine) as well as specific functions (like cysteine desulfuration, serine dehydratase) (Figs. 6 and 7). NH_4_^+^ may occupy one or two pockets and block the PLP binding as well as the R77 in-ward conformation (Fig. 4 and Fig. S4). Given that NH_4_^+^ is a possible catalytical product of the cysteine desulfuration catalyzed by psTD (Fig. 2D), this phenomenon might be a feedback inhibition of the enzymatic reaction (Figs. 3, 4, and 7E), which is confirmed by the in-gel activity assay (Fig. 3D). On the other hand, R77 with inward orientation can stabilize PLP binding through salt bridge interaction in PI, while R77 with outward orientation may open an access channel (PIII) of active site (Fig. 7). The mutation analysis shows R77E significantly changes the charge of gate residue as well as the flexibility of main-chain around residues 76-78, which leads to block small molecular traffics (Fig. 4). The MEP of active site and dynamics and flexibility of R77 may play a key role in enzyme activity, which is similar to the roles of R62 from hCGL and its structural dynamics described in previous report (Yan et al, 2017). R77, as the gate-keeper, is a potential switch to control the small molecules exchange at the gate and in the active site as well as the reaction type and process at active site (Fig. 7E). Overall, the above results shed light on mechanisms of psTD catalyzing cysteine to form H_2_S, which consists well with the results disclosed by genetic, proteomic and biochemical results in this study.

Altogether, we first show that psTD mediates cysteine desulfuration both *in vivo* and *in vitro*. Given the broad distribution of psTD homologs in different bacteria, we speculate that some threonine dehydratases have evolved a novel function towards cysteine desulfuration, which benefits the producer to utilize cysteine as a sulfur source for better adapting external environments.

## MATERIALS AND METHODS

### Strains, media, and chemicals

*Pseudomonas stutzeri* 273 was isolated from the sediment samples collected by RV *KEXUE* in the East China Sea in the year of 2014 (Wu et al, 2016). *P. stutzeri* 273 and its mutants were incubated in LB broth (10 g/L peptone, 10 g/L NaCl, 5 g/L yeast extract, pH 7.0) under vigorous agitation at the speed of 150 rpm at 28 °C. *Escherichia coli* DH5α was used as the host for plasmid construction. *E. coli* S17-1 was used as a vector donor in conjugation. *E. coli* BL21 was used for recombinant protein overproduction. *E. coli* DH5α, *E. coli* SY327, *E. coli* S17-1, and *E. coli* BL21 were grown in LB medium at 37 °C with shaking speed of 150 rpm. When necessary, antibiotics were used at the following final concentrations: 25 μg/mL chloramphenicol (Cm), 25 μg/mL gentamicin (Gm), 100 μg/mL ampicillin (Amp), and 100 μg/mL kanamycin (Kan) (Zhang et al, 2020).

### H_2_S production assay

To detect H_2_S production of *P. stutzeri* 273, bacterial cells were transferred into 5 ml of LB with the addition of L-cysteine in glass tubes (18*150 mm). Paper strips with lead acetate were affixed at the top of the tubes with rubber stoppers. After shaking incubation for 12 h, the paper strips were photographed to detect the presence of black lead sulfide precipitates, which is correlated to the production of H_2_S. The estimation was done by visually matching the darkness of the paper strips (Xia et al, 2017).

### Proteomic analysis

*P. stutzeri* 273 cells were incubated in LB and LB with 8 mmol/L L-cysteine until OD_600_ was 0.8. Then proteins were extracted, separated and identified using liquid chromatography-tandem mass spectrometry (LC-ESI-MS/MS) analysis. The bioinformatic analyses of protein annotation, functional classification, functional enrichment and cluster analyses were performed. The heat map of differently expressed proteins (1.5-fold change cutoff and *P* value less than 0.05) was made by the software Heml 1.0.3.3.

### Bioinformatics analysis

*P. stutzeri* 273 was collected in China General Microbiological Culture Collection Center under collection number CGMCC 7.265. The complete genome sequence of *P. stutzeri* 273 has been deposited at GenBank under the accession number CP015641 (Wu et al, 2017). The gene sequences of *threonine dehydratase* (*psTD*, accession number PS273GM_RS04065), *cystathionine* γ*-lyase1* (*CSE1*, accession number PS273GM_RS00775) and *cystathionine* γ*-lyase2* (*CSE2*, accession number PS273GM_RS06855) were obtained from GenBank. The consensus phylogenetic tree of psTD in *P. stutzeri* 273 with other related proteins obtained from GenBank was constructed by the maximum likelihood method with MEGA 7.0 (Kumar et al, 2016). Weblogo of amino acid residues according to the multiple sequence alignment with psTD and other proteins in TD family (MSA-TD) was created in Weblogo (http://weblogo.berkeley.edu/logo.cgi). The MSA-TD was mapped to the psTD structure using ConSurf Server (https://consurf.tau.ac.il/) (Ashkenazy et al, 2010). The proteins used in MSA-TD are shown in Supplementary, Fig. S3B.

### Construction of deletion mutants and complementation strains in *P. stutzeri* 273

Gene knockout in *P. stutzeri* 273 was made following the conjugation method described previously (Wu et al, 2017; Zheng et al, 2020). Fragments for mutant construction were amplified from the chromosome of *P. stutzeri* 273 by primers shown in Supplementary Table S1. Purified homologous fragments upstream and downstream of the target region were digested and ligated into the suicide vector pEX18Gm containing an *oriT* for conjugation. The constructed vector was transferred into *E. coli* SY327 and then *E. coli* S17-1 in turn. Using *E. coli* S17-1 as a donor strain, the constructed vector was transferred into *P. stutzeri* 273 by intergeneric conjugation at 28 °C for 48 h. After mating, cells were plated on LB agar plate with Cm and Gm to screen for single-event positive recombinant strains. The individual colony was used for second crossover. Sucrose counter-selection produced mutants without the pEX18Gm region. All double-recombination mutant candidates were verified by PCR amplification and sequencing. The primers used for validating complete removal of the target region from the host genome were shown in Supplementary Table S1.

The plasmid pUCP18 was used to construct complementary strains. The gene of *psTD* together with its native promoter was amplified from the wild-type *P. stutzeri* 273 by primers listed in Supplementary Table S1. The purified PCR product was inserted into *Hin*dIII/*Bam*HI site of pUCP18 to produce pUCP18-*TD*. Then pUCP18-*TD* was transferred into mutant strain Δ*TD*. The final complementary strain Δ*TD*/*cTD* was verified by PCR amplification and sequencing.

### In-gel detection of H_2_S producing enzyme

The enzymes degrading L-cysteine and forming H_2_S were detected using an in-gel activity assay with bismuth staining (Basic et al, 2017; Yoshida et al, 2010). Briefly, the cell lysates obtained by sonication were applied to Native-PAGE gel (12%). After electrophoresis, the gel was incubated in 100 mmol/L triethanolamine-HCl pH 7.6, 10 μmol/L pyridoxal 5-phosphate monohydrate, 1.0 mmol/L bismuth trichloride, 10 mmol/L EDTA and 20 mmol/L L-cysteine at 37 °C for 30 min or 3 h. H_2_S formed during the enzymatic reaction precipitated as insoluble bismuth sulfide, and H_2_S producing enzymes appeared as brown to black bands in the gels.

### Expression and purification of psTD or its mutants

The gene encoding psTD or its mutant was cloned into a pET-28a expression vector incorporating an N-terminal His tag fusion. The vector pET-28a containing psTD encoding gene was used as a template for construction of different mutants of psTD. The mutation in the 77^th^ amino acid (R77) of psTD was introduced by site-directed mutagenesis using KOD -Plus-Mutagenesis Kit (TOYOBO, Japan) to express psTD R77A (R is replaced with A), R77E (R is replaced with E), and R77K (R is replaced with K), respectively. The primers used for site-directed mutagenesis were shown in Supplementary Table S1. To express psTD or its mutants, plasmid containing different target gene was transformed into *E. coli* BL21 cells (Zheng et al, 2020). An overnight culture of *E. coli* BL21 cells containing different expression vector was inoculated into 1 L LB broth. Cultures were grown for 3 h at 37 °C with aeration (150 rpm) until an OD_600_ of 0.8 was reached. Then the expression was induced by 0.1 mmol/L IPTG at 16 °C for 12 h. Thereafter, cells were collected by centrifugation (8,000 × *g*, 20 min), resuspended in lysis buffer (50 mM Tris, 150 mM NaCl, pH 8.0) and subjected to sonication. Following lysis, the extract was centrifuged, filtered, and injected into a HisTrap high-performance (HP) column (5 mL) (GE Healthcare, America). The proteins were eluted with increasing concentrations of imidazole buffer (50 mM Tris, 150 mM NaCl, and 50-250 mM imidazole). After overnight dialysis to Tris buffer (10 mM Tris, 150 mM NaCl, 10% glycerol, pH 8.0), the protein solution was applied to a HiTrapTM Q HP column (GE Healthcare) and eluted with linear gradient of 0.15-2.0 M of NaCl in 10 mM Tris (pH 8.0). After overnight dialysis to Tris buffer (10 mM Tris, 150 mM NaCl, pH 8.0), active fractions were collected and concentrated by ultrafltration (MWCO 10 kDa, Millipore), and loaded onto a HiloadTM 16/600 superdexTM 200 column (GE Healthcare, USA) pre-equilibrated with 10 mM Tris (pH 8.0) containing 150 mM NaCl. Bound proteins were eluted with the equivalent buffer at a flow rate of 1 ml/min. The purity of fractions collected was determined by SDS-PAGE. Protein aliquots were quick-frozen in liquid nitrogen and stored in −80 °C till future use.

### Liquid chromatography-mass spectrometry (LC-MS) analysis

To find out the products from L-cysteine catalyzed by psTD, the mixture of 20 mmol/L L-cysteine and 0.1 mg/mL psTD in Tris buffer (50 g/L Tris, 150 g/L NaCl, pH 7.5) was incubated at 37 °C for 2 h. After protein precipitation by trichloroacetic acid, 1.75 mL borate buffer (1 mol/L, pH 9.0), 750 µL methanol, 1 mL enzymatic mixture, and 30 µL of diethyl ethoxymethylenemalonate (DEEMM) were mixed in a screw-cap test tube, treated in ultrasound bath over 30 min, and then heated at 70 °C for 2 h allow complete degradation of excess DEEMM and reagent byproducts (Gomez-Alonso et al, 2007). After derivatization, the reaction products were analyzed by liquid chromatography (Agilent Technologies)-mass spectrometry (BRUKER, maxis plus) (LC-MS). Pyruvate concentration in the enzymatic mixture was measured using 2,4-dinitrophenylhydrazine by monitoring the absorbance at 515 nm in relation to a standard curve (Anthon & Barrett, 2003).

### Crystallization and structural determination

Purified psTD or corresponding mutant was mixed with PLP (2 mmol/L) to a final concentration at 10.8 mg/mL (psTD or mutants) in the buffer containing 20 mmol/L Tris-HCl, 150 mmol/L NaCl, 1 mmol/L β-mercaptoethanol, pH 8.0. psTD, its corresponding mutant or psTD-PLP/psTD mutant-PLP complexes were screened for crystallization conditions using sitting-drop vapor diffusion at 6, 12 and 18[mg/mL. Crystals were formed in 1.8[mol/L ammonium sulfate, 0.1 mol/L Bis-Tris (pH 6.5), 2% (v/v) PEG 550 (E11 condition), or 0.1 mol/L NaAc·3H_2_O (pH 4.6), 3.5 mol/L HCOONa (C6 condition) within a week. Crystals were harvested and soaked in well solution containing 25% glycerol before flash frozen in liquid nitrogen. Crystals data were collected at 100 K at the beamline BL17U1 and BL19U1 at the Shanghai Synchrotron Radiation Facility (SSRF, China). The homolog model of TD was used for molecular replace with PHENIX, and the structure model was manually modified and refined by PHENIX through iterative cycles. The final refinement statistics are given in Table 1. All Figures were created with PyMOL (http://www.pymol.org/).

### Statistical analysis

All experiments were performed in triplicate and the data were expressed as mean ± standard deviation. The statistical analyses were performed with one-way analysis of variance (ANOVA). A multiple comparison Tukey test was used to evaluate if significant differences among treatments existed.

## Data availability

All proteomics related data have been deposited to the ProteomeXchangeConsortium via the PRIDE partner repository (dataset identifier PXD011469). The structures of psTD (C6 condition, native structure with PLP bound), psTD (E11 condition), psTD (psTD-PLP co-crystal complex with C6 condition), and psTD mutant (R77E mutation with C6 condition) have been deposited in the Protein Data Bank (PDB) under the accession codes 7DAP, 7DAQ, 7D8Y, and 7DAR, respectively.

### Acknowledgments

This work was funded by the China Ocean Mineral Resources R&D Association Grant (Grant No. DY135-B2-14), Strategic Priority Research Program of the Chinese Academy of Sciences (Grant No. XDA22050301), National Key R and D Program of China (Grant No. 2018YFC0310800), the Taishan Young Scholar Program of Shandong Province (tsqn20161051), Qingdao Innovation Leadership Program (Grant No. 18-1-2-7-zhc) for Chaomin Sun, and General Project of the National Natural Science Foundation of China (11179012), National Key Basic Research and Development Plan (973 Plan, 2011CB710800) for Wen Zhang.

## Author contributions

NM and CS conceived and designed the experiments. NM performed majority of the experiments and analyzed the data. NM, YS, and WZ purified proteins. YS, WZ design the crystallization experiments, grew crystals, collected data, solved the structures and carried out the structure analysis. NM, WZ and CS prepared the Figures and wrote the paper with all the inputs of all authors.

