## Supplemental Figures and Tables for "A novel deep-sea bacterial threonine dehydratase drives cysteine desulfuration and hydrogen sulfide production"

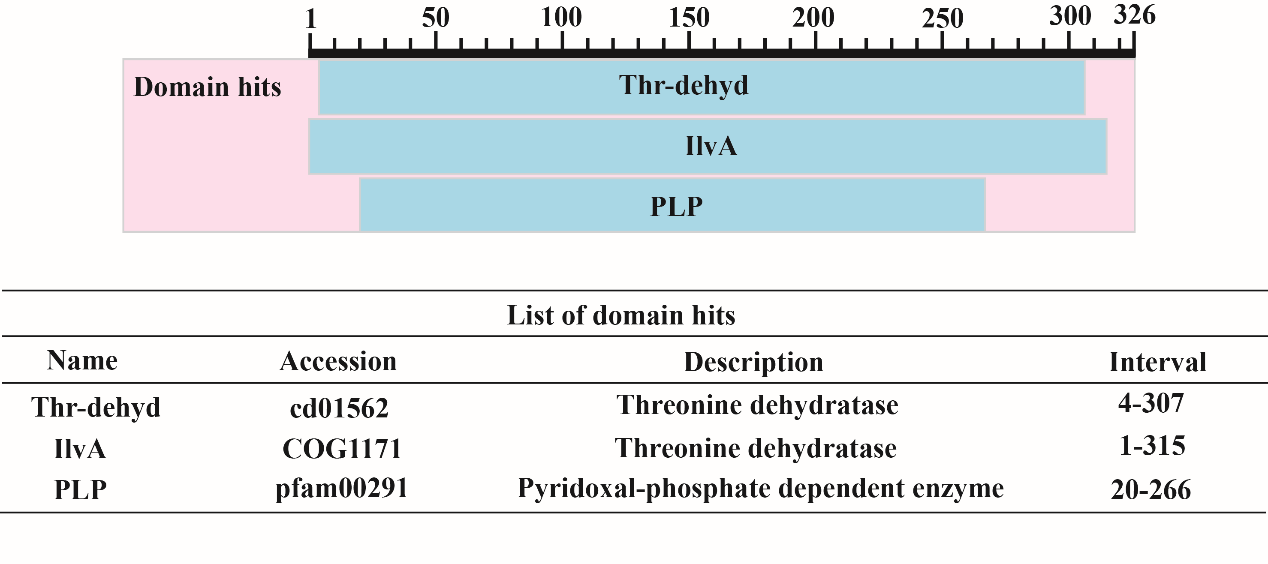

**Fig. S1** Typical domain hits existing in psTD.

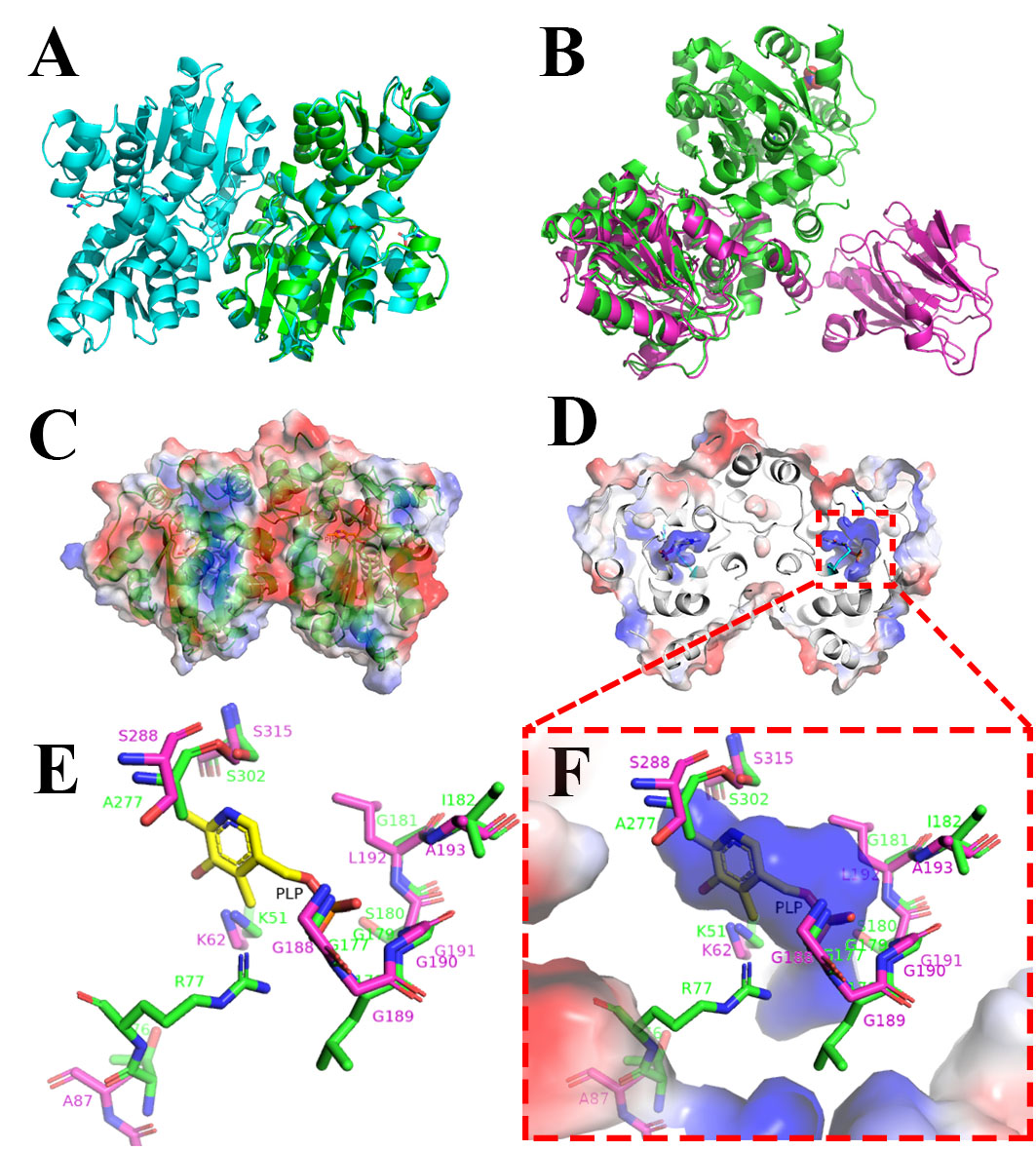

**Fig. S2 Crystal structure of psTD.** (**A**) Crystal structure of psTD asymmetric homodimer (PDB: 7D8Y). The structures are very similar between chain A (in cyan) and B (in green) (superpose RMSD=0.193). (**B**) The structure comparison by superpose 1TDJ (in purple) and psTD (in green): catalytic domains are conserved (RMSD=1.677) although their homology identity is only 31%. MEP for TD dimer: both outside surface (**C**) and site inside (**D**). (**E**) Key residues comparison around PLP between psTD (green carbon) and 1TDJ (purple carbon). (**F**) key residues and MEP for active site of psTD.

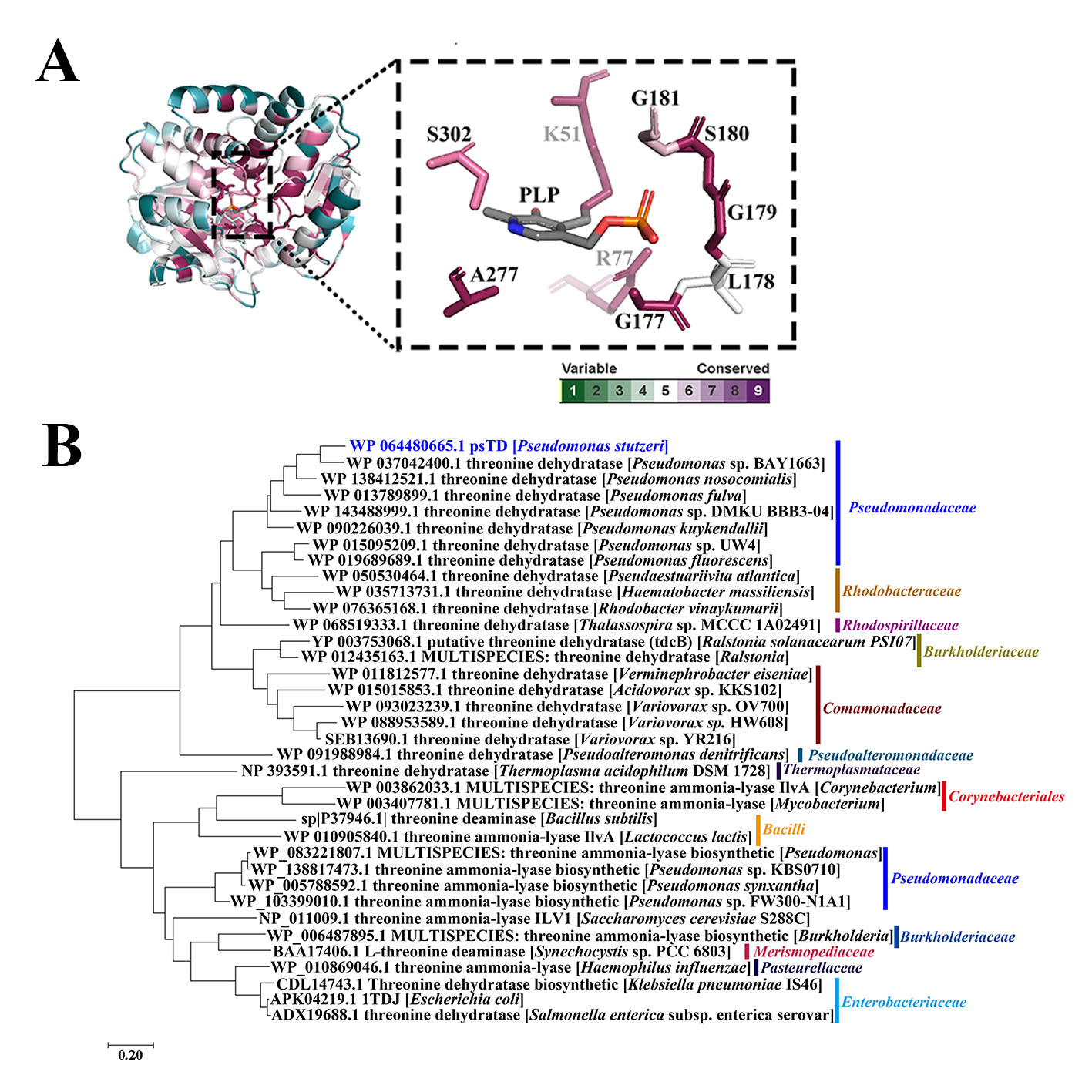

**Fig. S3 Sequence conservation of psTD.** (**A**) The sequence conservation mapping to the psTD structure (ConSurf view). Multiple sequence alignment was analyzed with TD and other proteins in threonine dehydrase family (MSA-TD). Then, the MSA-TD was mapped to the psTD structure using ConSurf Server (https://consurf.tau.ac.il/). The Figure was created using Pymol version 2. (**B**) The consensus phylogenetic tree of TD with other related strains according to MSA-TD by maximum likelihood method (accession numbers are indicated before the protein name).

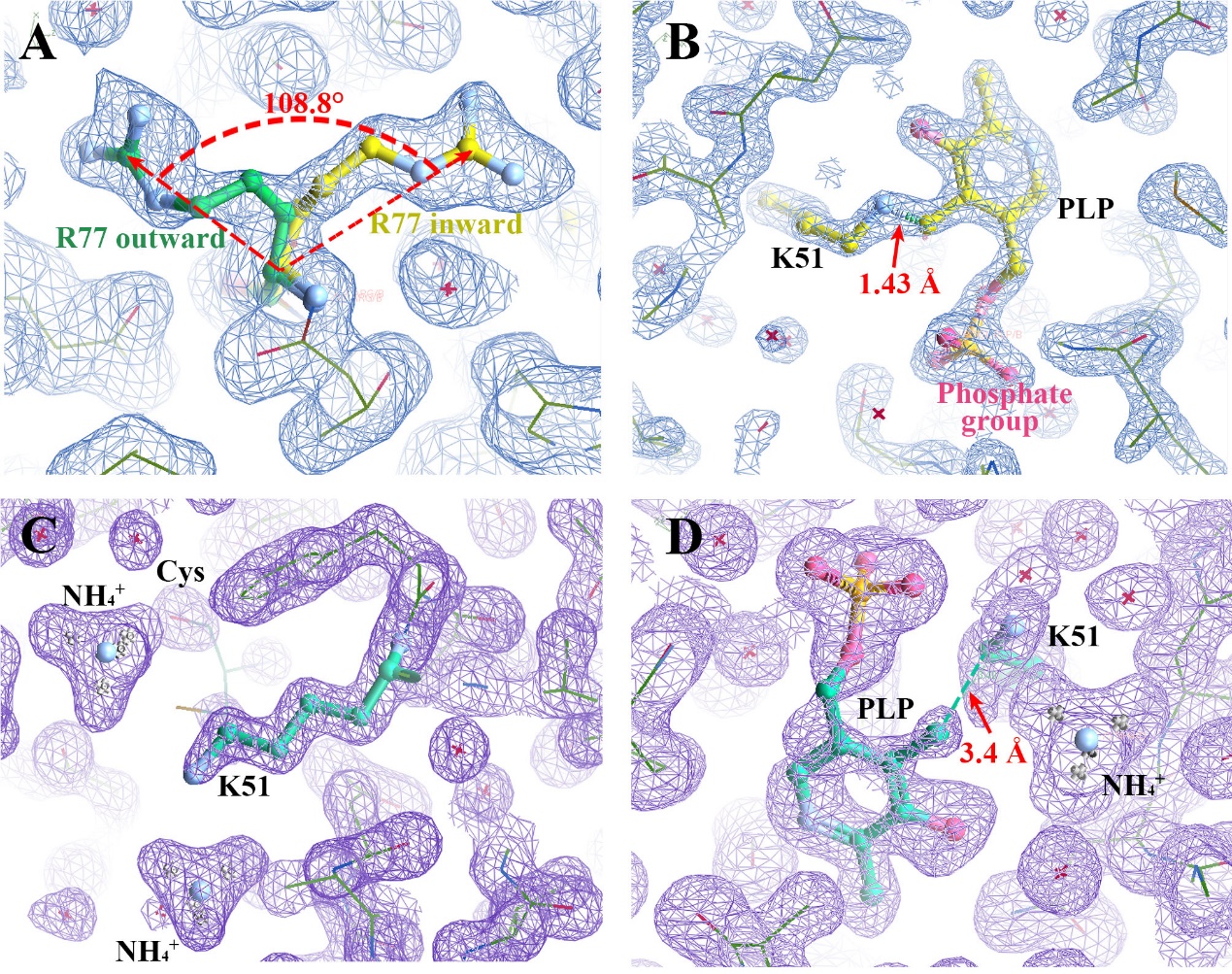

**Fig. S4 The crystal density map for the PLP and key residues K51 and R77.** (**A**) The alternative conformation of R77 (the angle between inward and outward orientation is about 108.8°, chain B of 7D8Y, map RMSD=0.6). (**B**) The PLP covalently bound to K51(bind length=1.43Å, chain B of 7D8Y, map RMSD=2.0). (**C**) K51 and two NH_4_^+^ in the active site (chain A of 7DAQ, map RMSD=1.7). (**D**) PLP bound to K51 with non-covalent bind (distance~3.4 Å, chain B of 7DAQ, map RMSD=0.7) and possible weak density for PLP at chain B of 7DAQ.

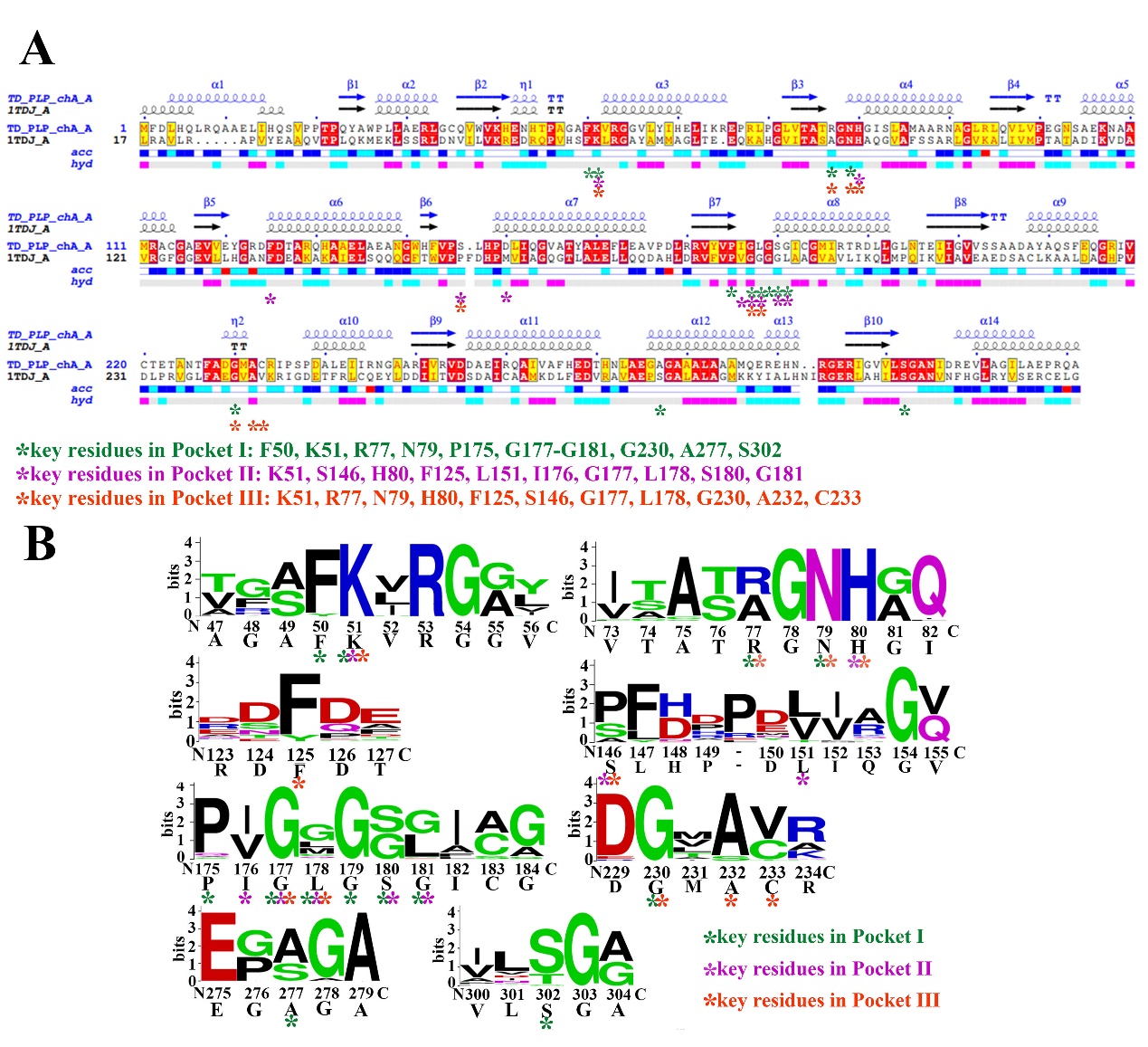

**Fig. S5 Key residues in Pockets I/II/III.** (**A**) Structure-based sequence alignment of psTD and 1DTJ. The sequences are directly extracted from Protein Data Bank (PDB) 1DTJ and psTD structure, and the sequence alignment was based on above two structures by ENDscript and ESPRIPT with default settings (http://espript.ibcp.fr/ESPript/ENDscript/index.php) (Robert & Gouet, 2014). Note: the key residues of Pocket I, Pocket II, and Pocket III in psTD are highlighted with different colored asterisks. (**B**) Weblogo of amino acid residues around Pocket I/II/III according to MSA-TD. The numbers are the locations of amino acids in TD. psTD sequence is underneath of the horizontal axis.

**Table S1** Primers used in gene knockout, mutant verification, gene complementary, TD encoding gene over-expression, and site-direct mutagenesis.

| Project | Primer name | Primer sequence |
| --- | --- | --- |
| Gene knockout | TD-L-F | CGCGGATCCGCGCGGCAGCCGCAACTGACGCC |
|  | TD-L-R | CGCCGTTCATCGAGTGAGGTCGACGGCTCCTCGGCGGGTT |
|  | TD-R-F | AACCCGCCGAGGAGCCGTCGACCTCACTCGATGAACGGCG |
|  | TD-R-R | CCGGAATTCCGG GCTGACCCAG GCCATGCACC |
|  | CSE1-L-F | CGCGGATCCGCGCTGTACAAAGCCAAGCGCAA |
|  | CSE1-L-R | TGTGGTTCTAAAGGAATAAGCTGTGTAATGGTACTGGCTG |
|  | CSE1-R-F | CAGCCAGTACCATTACACAGCTTATTCCTTTAGAACCACA |
|  | CSE1-R-R | CCGGAATTCCGGGGCAATTGGAATGTCGCTGAG |
|  | CSE2-L-F | CGCGGATCCGCGAGTAAGAGTTGCTAGCGCGT |
|  | CSE2-L-R | TTTGGTTTTAAGGTCCCTCGCATCCGGTATATGTGTGCCT |
|  | CSE2-R-F | AGGCACACATATACCGGATGCGAGGGACCTTAAAACCAAA |
|  | CSE2-R-R | CCGGAATTCCGGATTGCGCTCGGAGTTTCGTT |
| Mutant verification | TD-F | ATTGCCGTGCTGGGATTGA |
|  | TD-R | CAATGTGCAGTGCCTGGATG |
|  | CSE1-F | GCCAGACCAAGCTGATCGAA |
|  | CSE1-R | GGCTTATTTTGCAGCCGACC |
|  | CSE2-F | CTGTACGGCTTCGCTAGTCT |
|  | CSE2-R | GGCGAGTACGACATTCTCCT |
| Gene complementary | TD-C-F | CGCGGATCCGCGGATGTTCGATCTACACCAACT |
|  | TD-C-R | CCCAAGCTTGGGTCAGGCGGTCTGGCCAGCCG |
| TD over-expression | TD-P-F | CGCGGATCCGCG ATGTTCGATC TACACCAACT |
|  | TD-P-R | CCCAAGCTTGGG TCAGGCGGTCTGGCCAGCCG |
| site-direct mutagenesis | 77E-F | GAGGGCAATCACGGCATCAGCCTGGCGA |
|  | 77E-R | GGTTGCGGTGACGAGGCCCGGCAAG |
|  | 77A-F | GCTGGCAATCACGGCATCAGCCTGGCGA |
|  | 77A-R | GGTTGCGGTGACGAGGCCCGGCAAG |
|  | 77K-F | AAGGGCAATCACGGCATCAGCCTGGCGA |
|  | 77K-R | GGTTGCGGTGACGAGGCCCGGCAAG |

**Table S2** Measured Michaelis-Menten kinetic parameters of psTD (Km and Vm) with respect to L-cysteine.

| L-cysteine^a^ | | | |
| --- | --- | --- | --- |
|  | *K*_m_ (mmol/L) | *V*_m_^b^ (U/mg) | References |
| psTD | 14.40±1.78 | 3.08±0.25 | This work |
| smCSE | 87±25 | 0.68±0.20 | (Dunleavy et al, 2016) |
| Human CSE | 2.75 | 0.14 | (Sun et al, 2009) |
| *P. intermedia* CSE | 0.7 | 4.2 | (Yano et al, 2009) |

a: L-cysteine kinetic assay: 20 mmol/L L-cysteine was added to a preparation containing 0.5 mmol/L lead acetate and 0.1 mg/mL psTD in HEPES buffer (100 mmol/L, pH 7.5). The formation of PbS induced by H_2_S generation was monitored by measuring the absorbance at 390 nm. Runs were performed in triplicate and kinetic parameters were reported as mean ± standard deviation. For quantification of initial rates, a molar extinction coefficient for PbS of 5500 mol^-1^cm^-1^ was used. b: One unit corresponds to 1 μmol of product formed min^-1^.
